# WMW-A: Rank-based two-sample independent test for smallsample sizes through an auxiliary sample

**DOI:** 10.1101/2021.06.24.449844

**Authors:** Yin Guo, Limin Li

**Affiliations:** School of Mathematics and Statistics, Xi’an Jiaotong University, Xianning West 28, Xi’an, China

## Abstract

Two-sample independent test methods are widely used in case-control studies to identify significant changes or differences, for example, to identify key pathogenic genes by comparing the gene expression levels in normal and disease cells. However, due to the high cost of data collection or labelling, many studies face the small sample problem, for which the traditional two-sample test methods often lose power. We propose a novel rank-based nonparametric test method WMW-A for small sample problem by introducing a three-sample statistic through another auxiliary sample. By combining the case, control and auxiliary samples together, we construct a three-sample WMW-A statistic based on the gap between the average ranks of the case and control samples in the combined samples. By assuming that the auxiliary sample follows a mixed distribution of the case and control populations, we analyze the theoretical properties of the WMW-A statistic and approximate the theoretical power. The extensive simulation experiments and real applications on microarray gene expression data sets show the WMW-A test could significantly improve the test power for two-sample problem with small sample sizes, by either available unlabelled auxiliary data or generated auxiliary data.

## Introduction

Two-sample independent test methods are commonly used in case-control studies for testing whether the case population is significantly different from the control population. A two-sample test is performed on the data of two random samples independently obtained from two given populations, respectively. The purpose of the test is to determine whether there’s significant difference between the two populations. One example is that the two populations correspond to research subjects who have been treated with two different treatments (one of them can be a new drug or placebo). The goal is to assess whether one of the treatments typically yields a better response than the other. Another example is to identify differentially expressed genes by a two-sample test from microarray or RNA-Seq gene expression datasets with disease cells and normal cells. The differentially expressed genes can further be used to analyze the disease pathway.

There are two types of two-sample test methods: parametric and nonparametric methods. Parametric methods such as two-sample t-test assume the parametric family of distributions for the two populations and test whether the location parameters of the two distributions are the same. Two-sample t-test can be used when the underlying population distributions are normal. Other well-known parametric tests include t-testR method [7], which performs the t-test method on the ranked data, and Welch test method [26], which accounts for the two samples with unequal variances.

Nonparametric testing methods do not assume any particular parametric family of probability distributions for the two populations. Examples of nonparametric testing methods include Wold-Wolfowitz [25], Wilcoxon-Mann-Whitney statistic [16, 27] and Kolmgorov-Smirnov test [8]. Wilcoxon-Mann-Whitney test (WMW, also called Wilcoxon rank-sum test) is a commonly used rank-based hypothesis test, which constructs a statistic to compare the overall ranks of the two populations. The distribution of the WMW statistic under null hypothesis is often tabulated exactly for small sample sizes, but for sample sizes above 20, approximation using the normal distribution is fairly good.

In recent years, many scholars have put the two-sample test into concrete application environments and put forward many new ideas and methods. For example, [4, 5] studied the two-sample test problem for multivariate data and non-Euclidean data. By defining a similarity measure on the sample space, a two-sample test was performed based on the similarity graph. [4] further investigated the situation where the sample sizes of two samples were not equal. In particular, for the equality of two multivariate distributions, common methods are similar to [4, 5], which use the distance between observation points to construct geometric graph, like Friedman-Rafsky test [11], the cross-match test [21]. Also, there are other solutions such as [15] tested the multivariate distribution of complex data by constructing a regression framework. Another common application is the test of the equality of two high-dimensional distributions. The classic Hotelling *T*^2^ test loses effect in such cases. [1, 6] improved Hotelling *T*^2^ test to make it suitable for high-dimensional data. The new *T*_2_ criterion proposed by [23] can achieve higher power under certain conditions. In particular, [3, 12, 29, 30] proposed test methods for the equality of means of high-dimensional data. In addition, there are some other application scenarios, such as [14, 17, 22] combined binary classification in machine learning with two-sample testing, [18, 19] studied censored life cycle data analysis, and [10] considered the partially paired two-sample problem in the biology field.

Although many two-sample test methods have been designed to test the location difference, few methods can deal with the problem of small samples, for which the sample sizes are small. The small sample problem exists in many applications of biological and medical studies. For example, there may not be enough volunteers who would like to test a new treatment, which will cause the lack of case observations. During the analysis of gene expression data, the observations of the gene expression after treatment may not be enough. The work in [28] proposed a pooled component test (PCT) for testing the equality of two multivariate sample mean vectors for small sample size by summing squared componentwise t-statistic. However, it is a parametric test for multivariate samples. WMW test can be also performed for small sample sizes, but it suffers from a low power. A simple extreme sample is that if the observations are *x* = [1, 2], *y* = [5, 6] from the two samples, respectively, the rank sum statistic has five possible values 3, 4, 5, 6, 7 with probabilities being 1/6, 1/6, 2/6, 1/6, 1/6. Thus the two-sided p-value computed by WMW test is 0.3333, which is clearly not small enough to reject the null hypothesis and thus produces a type II error. One reason for the low power in WMW test is that the sample space under null hypothesis is very small due to the small sample sizes. It is still challenging to deal with the small sample problem in two-sample test.

Note that in real applications, although case and control may have very small sample sizes, there might be plenty of unlabelled observations, which is potentially useful for two-sample test. For example, in the field of biomedicine, it is often difficult to study the differential genes for cancer subtypes by gene expression data from limited number of confirmed cancer subtypes. However, there is a large amount of unlabeled cancer gene expression data, whose cancer subtypes are unknown. In social surveys, research on privacy issues may be involved, and faced with a situation where there are only small samples with the labels related to the privacy issue. The large amount of available unlabeled data could potentially increase the test power for small sample sizes.

In this work, we develop a novel rank-based statistic WMW-A for two-sample independent test by introducing an auxiliary sample with data values. Rather than combining and ranking two samples, we combine the two samples and the auxiliary sample together, and rank the three samples in an increasing order. We define a three-sample WMW-A statistic as the gap between the average rank of the first sample (case sample) and that of the second sample (control sample) in the three combined samples. Intuitively, a large difference of average ranks implies a large difference between the two populations. The introduction of the auxiliary sample could increase the sample space of the statistic under the null hypothesis, and has potential to improve the test power. Simulation studies indicate the improved power of the WMW-A test over the traditional WMW test and t-test under different scenarios. The WMW-A test is also applied to several gene expression datasets for identifying differentially expressed genes between case and control observations, and the results indicate that the WMW-A test could obtain higher powers than the traditional WMW and t-test, with an acceptable type I error rate.

## Problem Statement and Notations

Suppose we have independent samples *X*_1_, · · ·, *X_m_* and *Y*_1_, · · ·, *Y_n_* from two populations *X* and *Y* with cumulative distribution functions (CDF) *F_X_* (*α_x_*) and *F_Y_* = *F_X_* (*α_x_* + *δ*), which are symmetric with *α_x_* and *α_y_*, respectively. Let **x** = [*x*_1_, · · ·, *x_m_*] be the data values associated with a random sample drawn from the first population *X*, and let **y**= [*y*_1_, · · ·, *y_n_*] be the data values associated with a random sample drawn from the second population *Y*. It is of interest to test the null hypothesis: *H*_0_: *F_X_* = *F_Y_* versus the two-sided alternative hypothesis: *H*_1_: *F_X_* ≠ *F_Y_*. We consider the small sample problem, which means the sample size sizes *m* and *n* are both small or either one is small.

### Mann–Whitney–Wilcoxon(WMW) test

Mann–Whitney–Wilcoxon test, also called Wilcoxon rank sum test, is a widely used nonparametric two-sample hypothesis testing method. The two random samples *X*_1_, · · ·, *X_m_* and *Y*_1_, · · ·, *Y_n_* are first combined together and ranked in an increasing order from 1 to *m* + *n*. If ties are present, the average rank is given to the tied values. Let *R_xi_* and *R_yj_* be the ranks of *X_i_* and *Y_j_* in the combined sample, and the WMW statistic is defined as

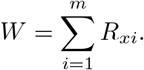

Under the null hypothesis, a mixture of low, medium and high ranks in each sample is expected, while under the alternative hypothesis one could expect lower ranks to dominate in one population and higher ranks in the other. The expectation and variance of the statistic under the null hypothesis *H*_0_ are

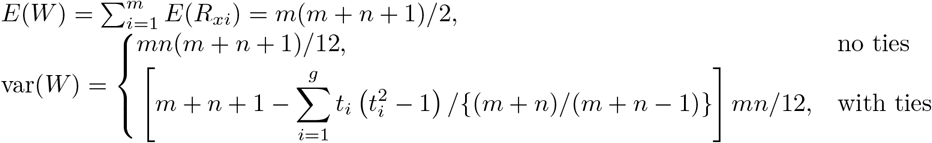

where *t_i_* is the number of observations in *i*-th tie, and *g* is the number of ties. When ties are present, normal or chi-square approximation for the statistic is often used to determine the p-value. For the case of small samples and no ties, exact permutation can be used to obtain the null distribution of the WMW statistic. Note that for small samples, for example *m* = *n* = 2, the least p-value can be obtained by WMW test is 0.333, thus the null hypothesis is never rejected. The small null probability space can lead to low power of WMW test for the case of small sample size.

## WMW-A test

To increase the power of two-sample test with small sample sizes, we introduce an auxiliary sample, which corresponds to unlabelled observations in real applications, and propose a nonparametric three-sample statistic. Theoretical properties of the statistic are also analyzed and given.

### The auxiliary population and sample

To deal with the small sample problem in two-sample test, we develop a novel nonparametric hypothesis test based on a WMW statistic by introducing an auxiliary sample from an auxiliary population. The test is called WMW-A test, where ‘A’ is for ‘auxiliary’.

We assume the auxiliary population *Z* by a mixed distribution of *X* and *Y*, with a cumulative distribution function

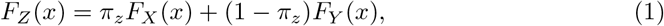

where *π_z_* is the prior probability that *Z* is from the population *X*, and 1 − *π_z_* is the prior probability that *Z* is from the population *Y*. *Z*_1_, · · ·, *Z_l_* is an auxiliary sample from the population *Z*. Since *Z* follows a mixture distribution, the member of the sample *Z_i_* is either from *X* or from *Y*, with prior probability *π_z_* and 1 − *π_z_*, respectively. The data values of the auxiliary sample are denoted as **z** = [*z*_1_, · · ·, *z_l_*], which are the large amount of unlabelled observations in real applications. For example, *X*-sample and *Y* -sample correspond to case and control observations, respectively, while the auxiliary *Z*-sample corresponds to the unlabelled observations. Note that in real applications, although case and control may have very small sample sizes, there might be plenty of unlabelled observations. Since it is unknown whether the unlabelled observations are from case population or control population, the data values can not be directly used in the traditional two-sample t-test or WMW test. We will develop a novel WMW-A test by making use of the unlabelled auxiliary observations to improve the test power.

### WMW-A statistic

Since the auxiliary *Z* is assumed to follow a mixture distribution of *X* and *Y*, the null hypothesis *H*_0_: *F_X_* = *F_Y_* is equivalent to 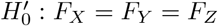. We first combine the three samples *X*_1_, · · ·, *X_m_*, *Y*_1_, · · ·, *Y_n_*, *Z*_1_, · · ·, *Z_l_* together and rank them in an increasing order. Let 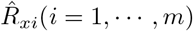 and 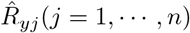 be the ranks of *X*_1_, · · ·, *X_m_* and *Y*_1_, · · ·, *Y_n_*, respectively, in the pooled sample of *X, Y* and *Z*. We define the WMW-A statistic as follows

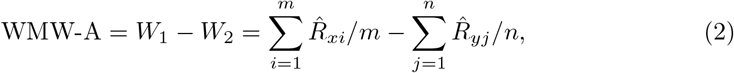

where the first term 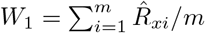 represents the average rank of the *X*-sample, and the second term 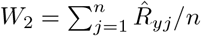 represents the average rank of the *Y* -sample, in the pooled sample of *X, Y* and *Z*. The main idea of the WMW-A test is shown in Figure 1. The WMW-A statistic is defined by the gap between the average ranks of the two samples in the large pooled sample, which is used to determine the strength of evidence against the null hypothesis. If the WMW-A statistic is large, it means that the locations of *X*-sample and *Y* -sample in the pooled sample are far from each other, and the null hypothesis tends to be rejected. The advantage of WMW-A statistic is that the rank is computed in a large pooled sample, which will produce more possibilities under the null hypothesis and extend the null probability space.

**Figure 1.**
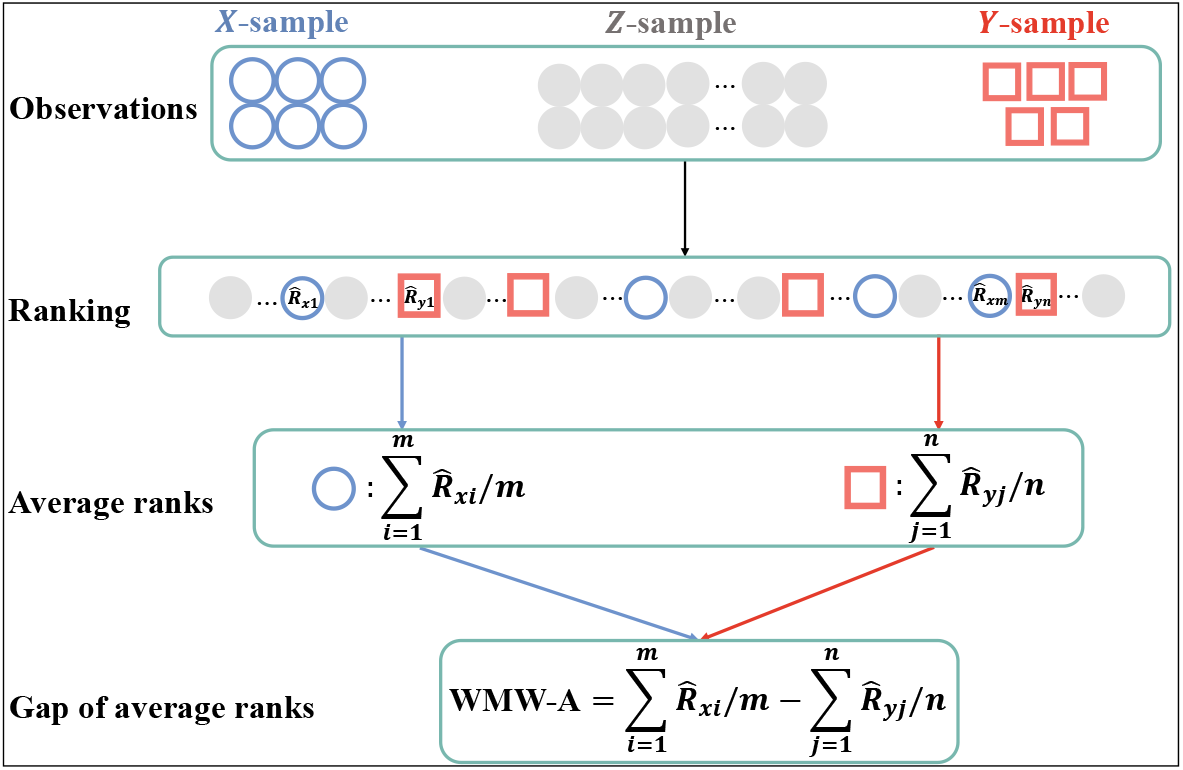
Main idea of WMW-A test.

### Properties of WMW-A statistic

We discuss the theoretical properties of the WMW-A statistic under the null hypothesis and alternative hypothesis, and give the following theorems.

#### Theorem 1

Under the null hypothesis, the WMW-A statistic follows a symmetrical distribution with the symmetrical center 0. Furthermore, under the null hypothesis, the mean and variance of WMW-A statistic, denoted by *μ* and *σ*^2^, are

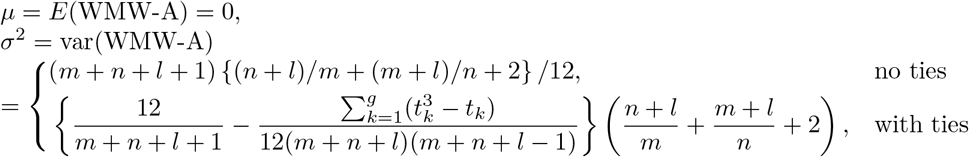

where *t_k_* represents the number of observations in the *k*-th tie, and *g* is the number of ties.

Proof: We first prove that the WMW-A statistic follows a symmetrical distribution under the null hypothesis.

Let *N* denotes the pooled sample size for convenience, i.e. *N* = *m* + *n* + *l*. Without loss of generality, suppose the ranks of *m* + *n* observations are taken out from the integer set of 1, …, *N*, which are *a*_1_, …, *a_m_, b*_1_, …, *b_n_*. Then the WMW-A statistic is denoted as

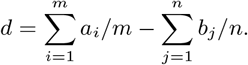

By introducing

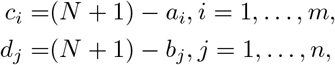

where 1 ≤ *c_i_, d_j_* ≤ *N, i* = 1, …, *m*; *j* = 1, …, *n*, we can obtain that

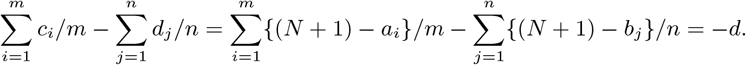

Thus it is easy to get the equations

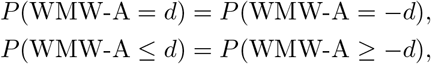

for *d* = −(*m* + *n* + 2*l*)/2, …, (*m* + *n* + 2*l*)/2, which means that the WMW-A statistic follows a symmetrical distribution, and the symmetrical center of the WMW-A statistic is the midpoint of −(*m* + *n* + 2*l*)/2, …, (*m* + *n* + 2*l*)/2, i.e.

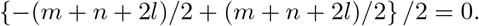

We next prove the mean and variance of the WMW-A statistic. The mean of the WMW-A statistic under the null hypothesis is

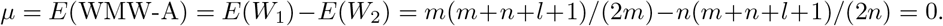

As for the variance of the WMW-A statistic, when there are no ties,

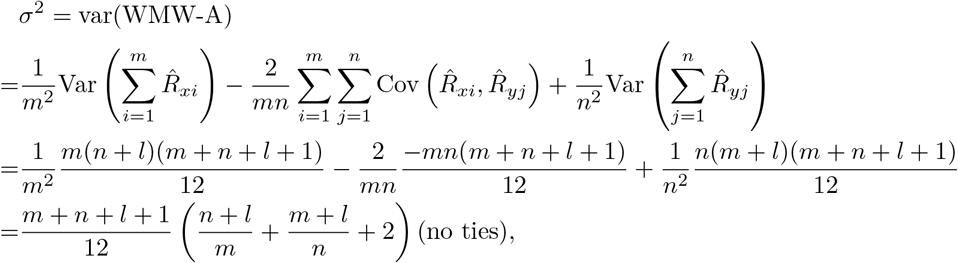

For the case with ties, the variance is

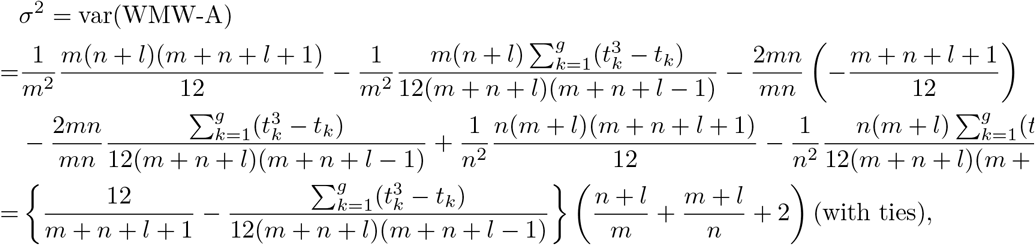

where *t_k_* represents the number of observations in *k*-th tie, and *g* is the number of ties.

To further analyze the properties of the WMW-A statistic under alternative hypothesis, we introduce two other populations *U* and *V*, where *U* ~ *F_U_* is a mixed distribution of *Y* and *Z*, and *V* ~ *F_V_* is a mixed distribution of *X* and *Z*. Then *Y*_1_, · · ·, *Y_n_, Z*_1_, · · ·, *Z_l_* can be considered as a sample with sample size *n* + *l* from population *U*, and *X*_1_, · · ·, *X_m_, Z*_1_, · · ·, *Z_l_* is a sample with sample size *m* + *l* from population *V*. *F_X_, F_Y_, F_U_* and *F_V_* are all unknown, but the empirical distributions could be estimated by the corresponding samples. We have the following theorem.

#### Theorem 2

Under the alternative hypothesis, the mean and approximated variance of the WMW-A statistic, denoted by 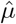 and 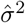, are

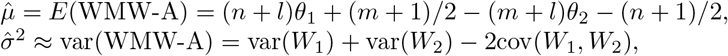

where

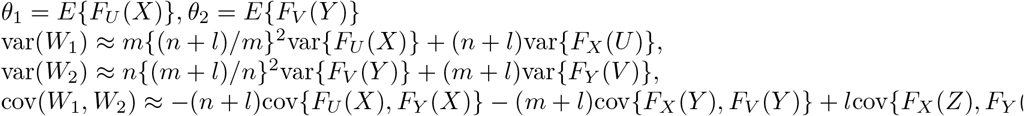

Proof: To obtain the mean of the statistic WMW-A, we take use of the relationship between WMW statistics and Mann–Whitney U statistics, that is, the statistics *W*_1_ and *W*_2_ can be written as:

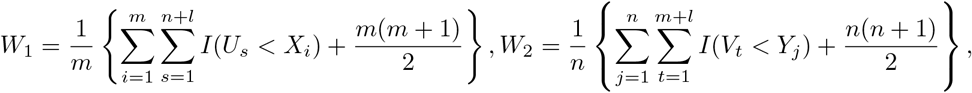

where *I*(·) is the indicator function whose values are either 1 or 0, and 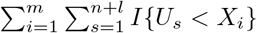 and 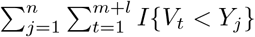 are the Mann–Whitney U statistics. The mean of the WMW-A statistic can thus be represented as

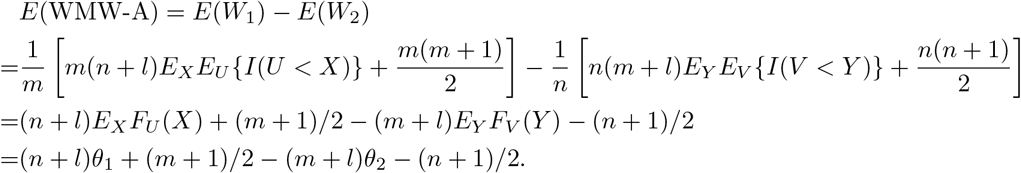

The expectation holds under both null and alternative hypothesis. Under the null hypothesis, *θ*_1_ = *θ*_2_ = 1/2, and thus the mean of the WMW-A statistic *μ* = 0, which is consistent with Theorem 1. Under the alternative hypothesis, the expectation, denoted by 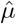, can be represented as (*n* + *l*)*θ*_1_ + (*m* + 1)/2 − (*m* + *l*)*θ*_2_ − (*n* + 1)/2, where *θ*_1_ and *θ*_2_ are not 1/2.

To derive the variance of the statistic WMW-A, we represent *W*_1_ and *W*_2_ in the form of asymptotic linear expansion. As a special case of the general results in section 13.4 of [24], the statistic *W*_1_ has the asymptotic linear expansion under alternative hypothesis

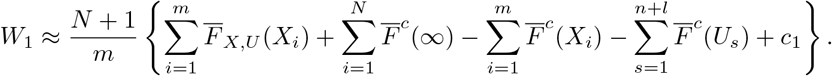

where *N* = *m* + *n* + *l*, 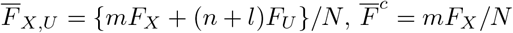, and *c*_1_ is a constant. Then the above can be further simplified as

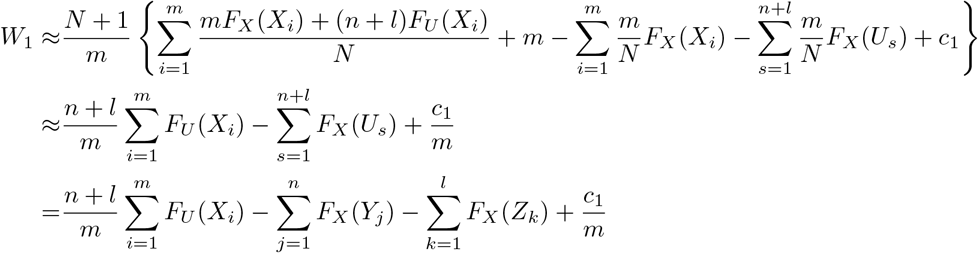

Similarly,

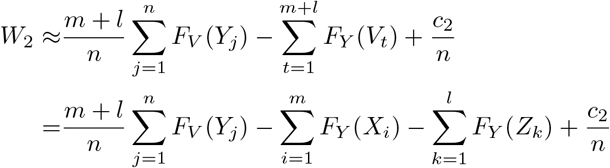

The variance of the statistic *W*_1_ under the alternative hypothesis thus can be approximated as

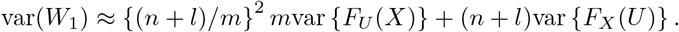

Similarly, we can obtain the approximated variance of the statistic *W*_2_:

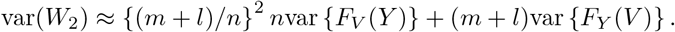

Such that the approximated variance of statistic WMW-A under alternatives is

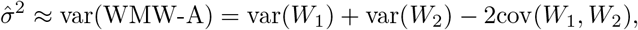

where

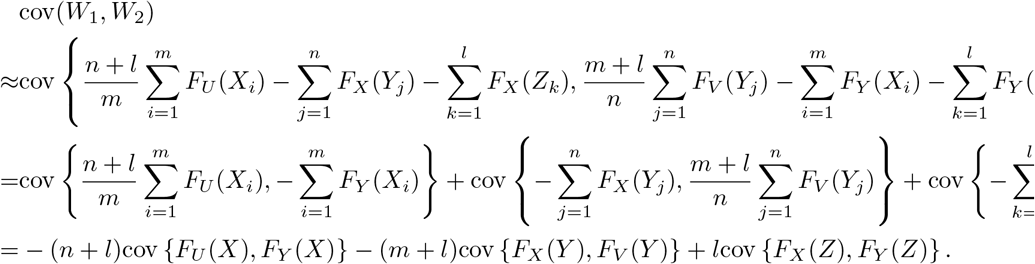

The cumulative distribution functions *F_X_*, *F_Y_*, *F_U_*, *F_V_* can be estimated by empirical distribution functions.

Through the above theorems and the laws of large numbers, we can get the theoretical asymptotic power of the WMW-A statistic in Theorem 3.

a. Under the null hypothesis, the WMW-A statistic follows a normal distribution asymptotically as *m* and *n* go to infinity:

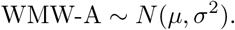
b. Under the alternative hypothesis, the asymptotic distribution of the WMW-A statistic as *m* and *n* go to infinity can be written as:

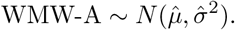

#### Theorem 3

The type II error rate for the two-sided WMW-A test as *m* and *n* go to infinity can be estimated by

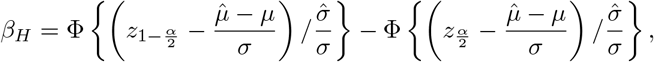

where *μ* and *σ* are the mean and standard deviation of the WMW-A statistic under the null hypothesis, 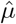 and 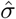 are the mean and standard deviation of the WMW-A statistic under the alternative hypothesis, Φ and *z*_1−*α*/2_ are the probability density function and the 1 − *α*/2 quantile of the standard normal distribution, respectively. The power of the WMW-A test thus can be estimated as 1 − *β_H_*.

Proof: For the two-sided hypothesis test, the null hypothesis is rejected if

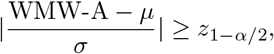

where *z*_1−*α*/2_ is the 1 − *α*/2 quantile of standard normal distribution. When the alternative hypothesis is true, since

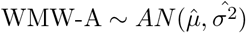

thus we have

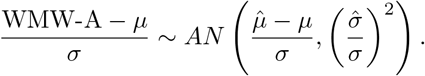

The type II error rate can be calculated by

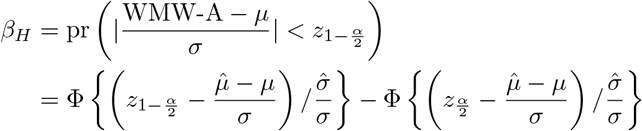

Thus, the power of the statistic is 1 − *β_H_*.

With large total sample sizes, the WMW-A statistic asymptotically follows normal distribution *N* (*μ, σ*^2^) under null hypothesis. Figure 2a shows the theoretical null distribution curves for an example with large sample sizes *m* = 50, *n* = 100 and *l* = 150 from *N* (0, 1) and *N* (1, 1). The vertical line shows the observed WMW-A statistics for observations **x** and **y**, and the test clearly rejects the null hypotheses.

**Figure 2.**
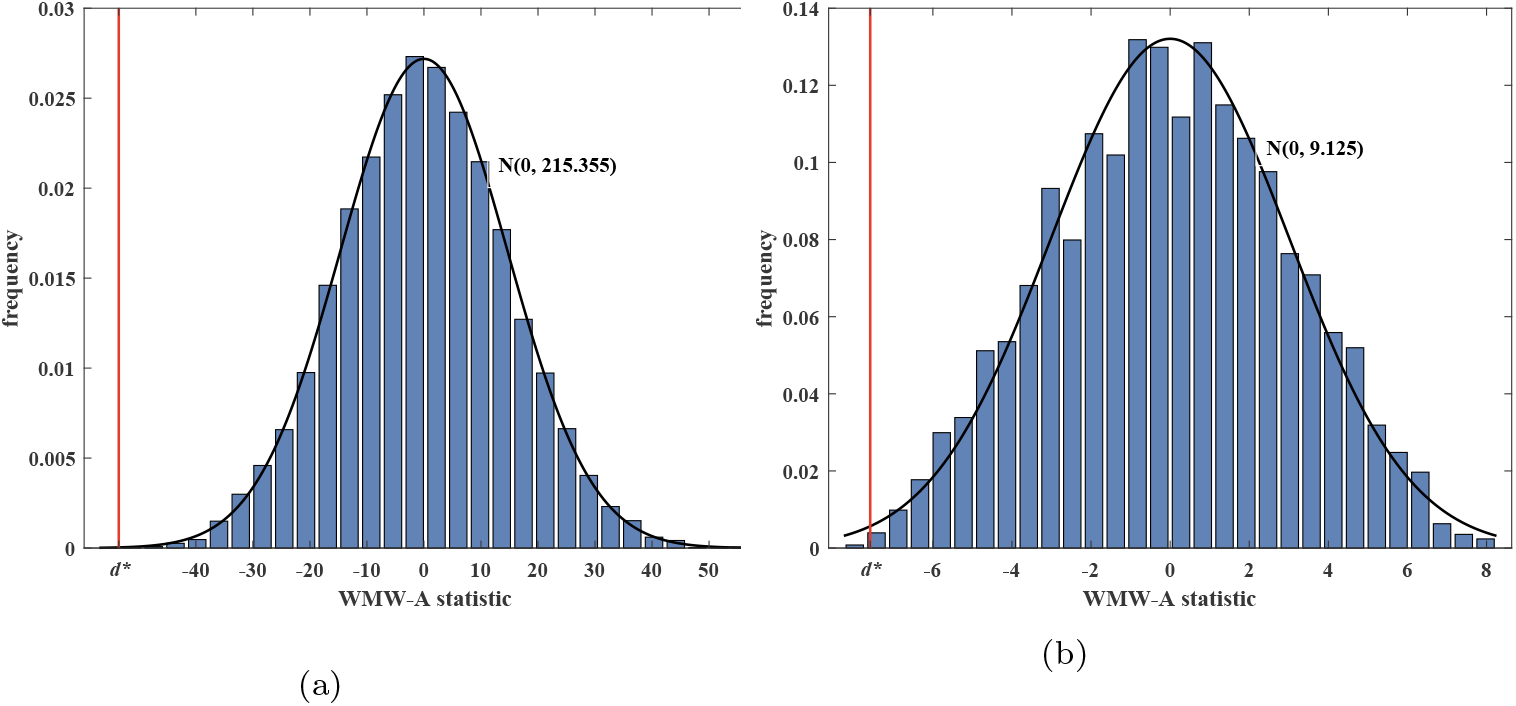
Frequency distribution histogram of WMW-A statistic by permutation and the theoretical asymptotic density function curve of WMW-A statistic, where the vertical lines represent the data values *d*^∗^ of WMW-A statistic computed by the given data observations. (a) large sample size, where *m* = 50, *n* = 100, *l* = 150, and **x** and **y** are generated from *N* (0, 1) and *N* (1, 1), respectively. (b) small sample size, where **x** = [1, 2], **y** = [7, 8, 9] and **z** = [2, 3, 4, 6, 7, 8].

### The WMW-A permutation test

The asymptotic normal property of the WMW-A statistic may not hold for small sample sizes, for example, *m* = 2, *n* = 3 and *l* = 6 in Figure 2(b). We thus propose a permutation strategy to obtain the empirical null distribution for the WMW-A statistic. Since the null hypothesis is 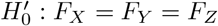, we permute the pooled samples from *X, Y* and *Z*, and then calculate the WMW-A statistic. The permutation procedure is performed for many times, and the null distribution for the WMW-A statistic can thus be obtained. The significance p-value can be finally computed based on the null distribution and the observed values of the WMW-A statistic. Note that to avoid outliers and achieve more stable results, we remove the observations of *Z*-sample outside the range of the observations of the pooled *X* and *Y* samples. We summarise the WMW-A permutation test in the algorithm box^1^. Figure 2 further shows the empirical null histograms of the WMW-A statistic by the permutation procedure. We can see that for the large sample sizes in Figure 2(a), the theoretical asymptotic null distribution is consistent with the empirical null distribution, and for the small sample sizes in Figure 2(b), the asymptotic normal property becomes weak.

**Table 1.**
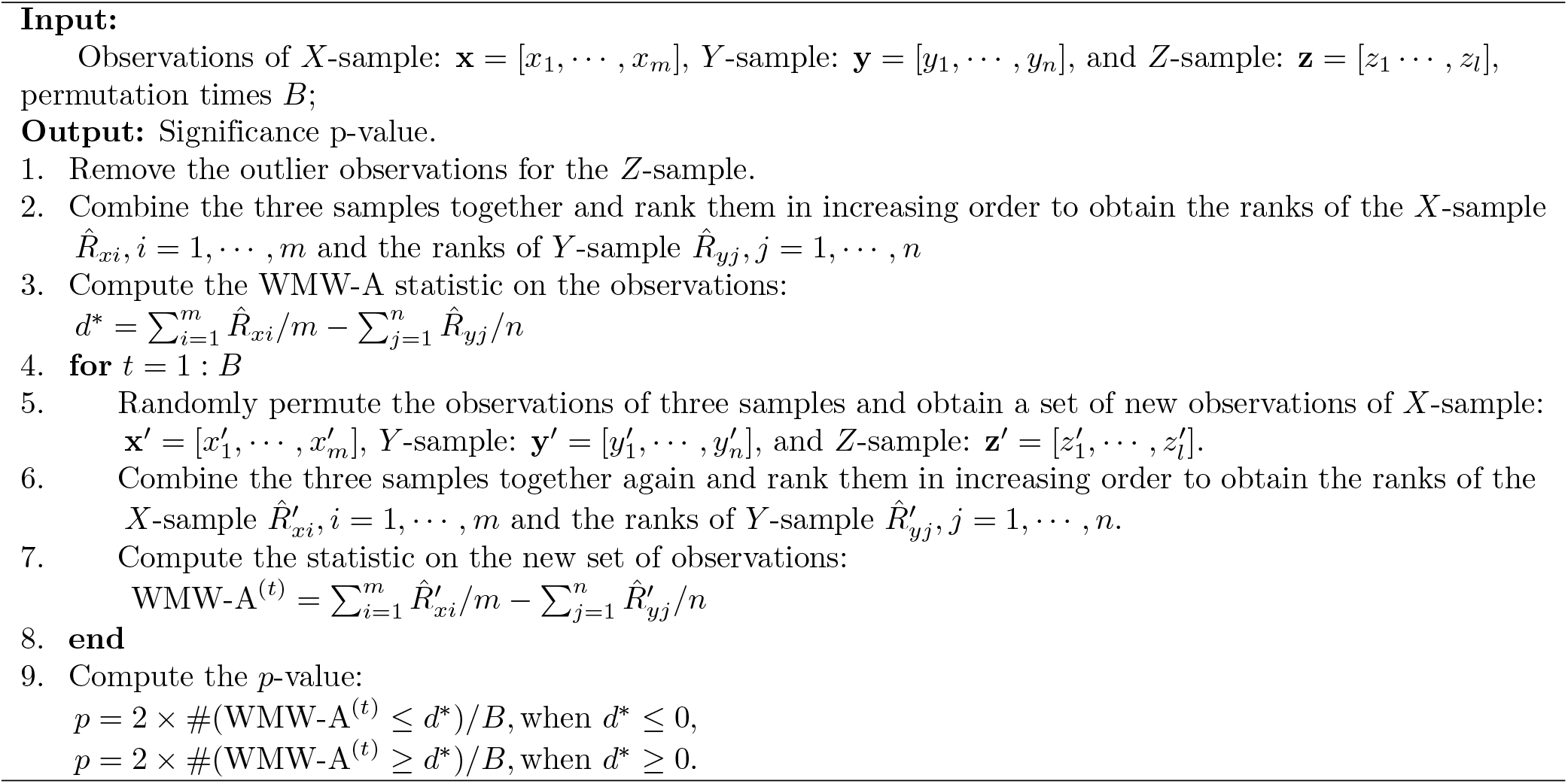
Algorithm: WMW-A permutation test.

## Simulation studies

### Data simulation

In simulation study, we choose *F_X_* and *F_Y_* to be normal distributions or Gamma distributions with location parameters *α_x_* and *α_y_*, respectively, to sufficiently evaluate the performance of the WMW-A test. The null hypothesis of the test is that the distributions *F_X_* and *F_Y_* have the same location parameter *α*, i.e. *H*_0_: *α_x_* = *α_y_*. We first generated *m* case observations **x** = [*x*_1_, · · ·, *x_m_*] for the *X*-sample and *n* control observations **y**= [*y*_1_, · · ·, *y_n_*] for the *Y* -sample from distributions *F_X_* and *F_Y_* (either the same or different), respectively. To perform the WMW-A test, we also generated *l* auxiliary observations **z** = [*z*_1_, · · ·, *z_l_*] for the *Z*-sample from the mixed probability in Equation (1) with parameter *π_z_*. The parameters for simulated data and parameters for the WMW-A test are shown in Table 2.

**Table 2.**
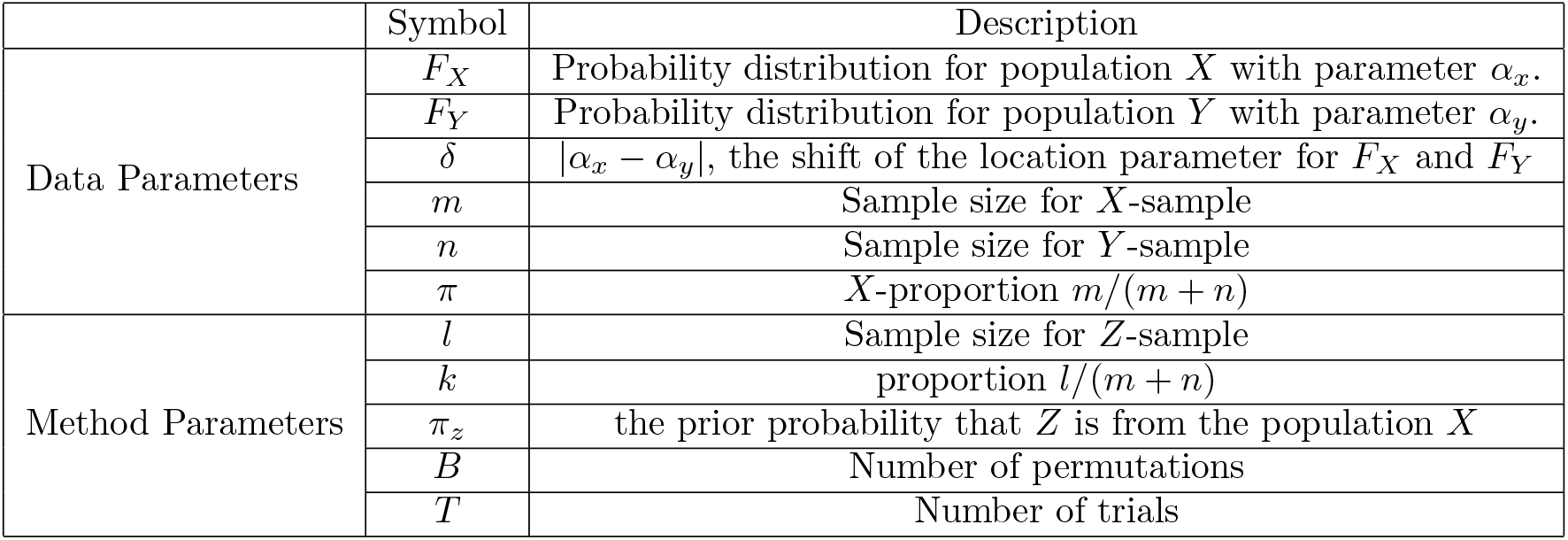
Notations and corresponding descriptions.

We compared the performance of the WMW-A test with classical t-test, t-testR [7], the WMW test and Welch test [26] methods on simulation datasets with different parameters. The power = *P* (reject *H*_0_|*H*_0_ is false) and type I error rate = *P* (reject *H*_0_|*H*_0_ is true) are estimated based on *T* Monte Carlo replicates.

### Experimental setting

To sufficiently evaluate the performance of the WMW-A test for the case of small sample, we studied the powers of the WMW-A test by considering different datasets with different properties as follows

a. *δ*, the degree of the shift between the two population distributions *F_X_* and *F_Y_*.
b. *m* and *n*, sample sizes of *X* and *Y*, and the proportion *π* = *m*/(*m* + *n*).

We take two types of population distributions as examples, Normal and Gamma distributions. For Normal distributions, we choose *F_X_* = *N* (0, 1), and *F_Y_* = *N* (*δ,* 1). For Gamma distributions, we choose *F_X_* = Γ(0.5, 1) and *F_Y_* = Γ(0.5 + *δ,* 1). The shift parameter *δ* is chosen from the set {0.5, 1, 1.5, 2}. We consider two cases for sample sizes *m* and *n*. One case is *m* = *n*, which are chosen from {2, 4, 5, 8, 10, 12, 15, 18, 20, 25}, and the other case is that we fix *m* = 5, and choose *n* from the set {2, 4, 8, 10, 15, 20, 25, 30, 40, 50}.

We also studied how the selection of *Z* influences the performance of WMW-A test. There are two parameters for selecting *Z*. One parameter is the sample size, *l*, and the other is the proportion of *Z* from the population *X*, *π_z_*. In our experiments, we take *l* = *k*(*m* + *n*), where *k* is chosen from the set {2, 5, 10}, and take *π_z_* from the set {0, 0.1, · · ·, 0.9, 1}. The number of permutations and trials are chosen as *B* = 2000 and *T* = 1000, respectively.

### Experimental results

We first evaluated our WMW-A method by the test powers. Figure 3 reported the test powers obtained by different test methods on different simulation datasets generated by Normal distributions with different parameters *δ*, *m*, *n* and *π*. We fixed *π_z_* = *π* and *k* = 5 for selecting the *Z* sample, and reported the experimental results on datasets generated by Gamma distributions in the Figure 4.

**Figure 3.**
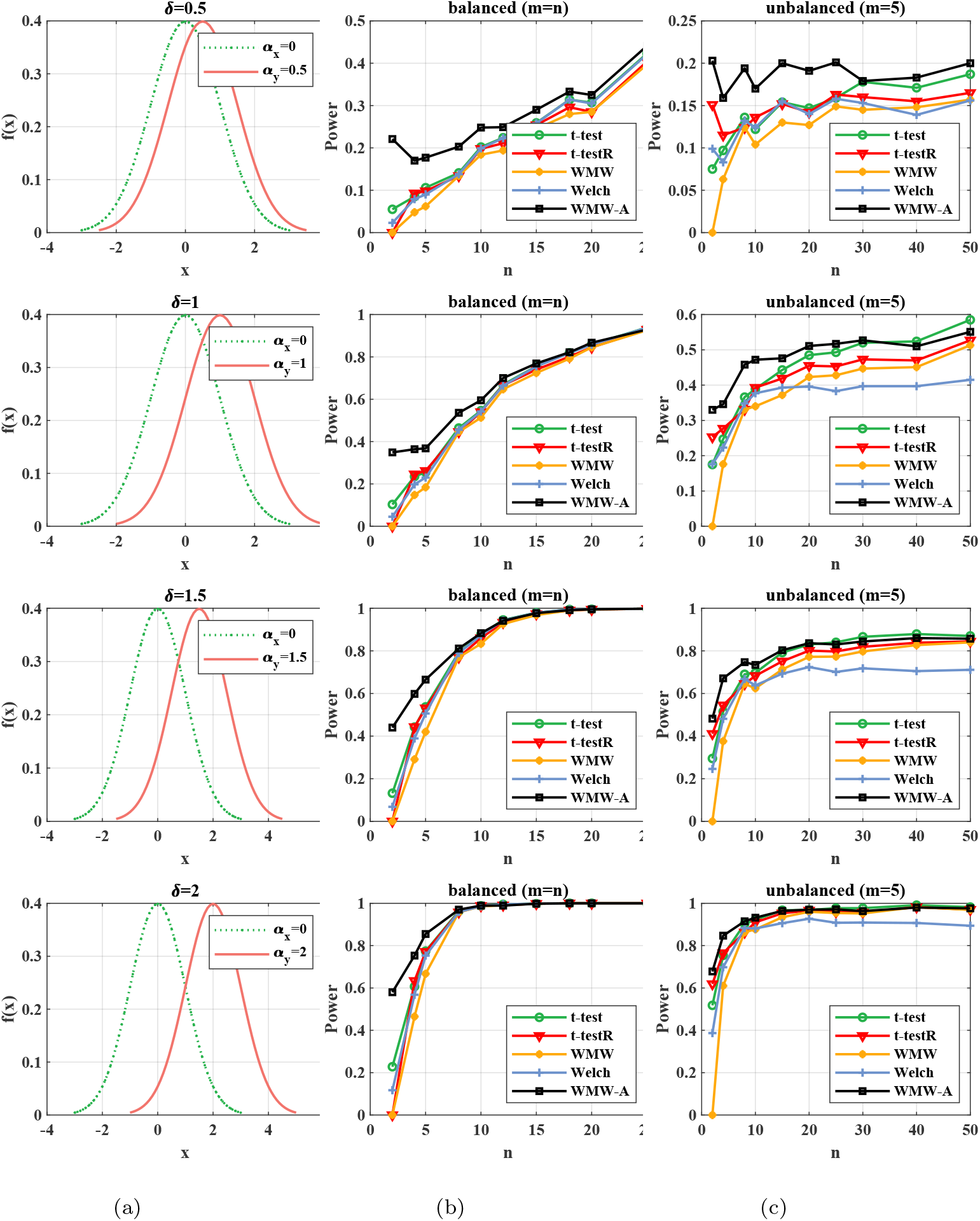
Powers of t-test, t-testR, WMW test, Welch test and WMW-A test (*k* = 5) for datasets generated from normal population distributions. (a).The distributions of the two populations *X* and *Y*, where the degrees of the shift between them are *δ* = {0.5, 1, 1.5, 2}, respectively, from the top to bottom; (b).Powers for the balanced case with increasing *m* = *n* for the corresponding *δ*; (c).Powers for the unbalanced case with fixed *m* = 5 and increasing *n* for the corresponding *δ*.

**Figure 4.**
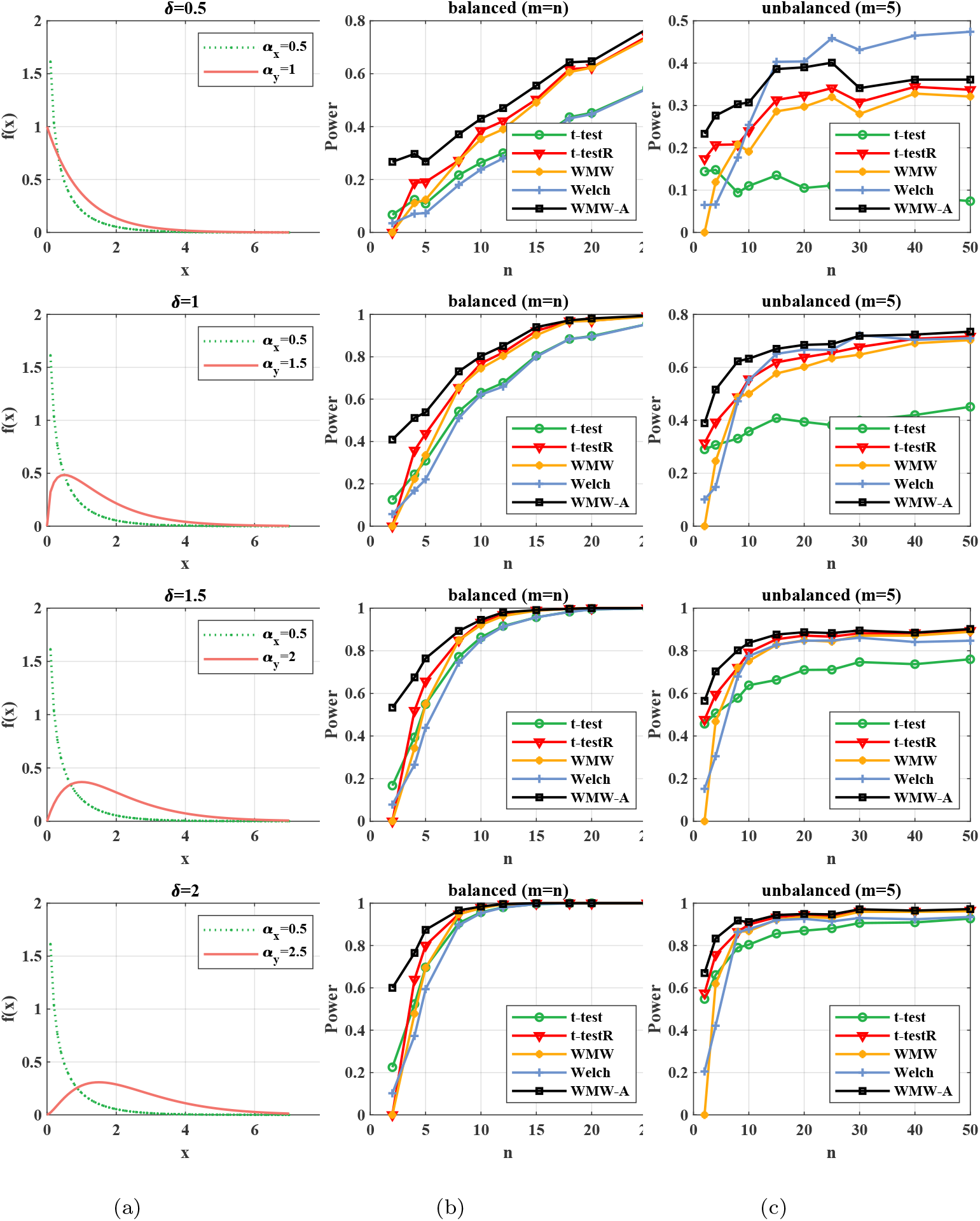
Powers of t-test, t-testR, WMW test, Welch test and WMW-A test (*k* = 5) for datasets generated from Gamma population distributions. (a).The distributions of the two populations *X* and *Y*, where the degrees of the shift between them are *δ* = {0.5, 1, 1.5, 2}, respectively, from the top to bottom; (b).Powers for the balanced case with increasing *m* = *n* for the corresponding *δ*; (c).Powers for the unbalanced case with fixed *m* = 5 and increasing *n* for the corresponding *δ*.

In Figure 3 (and Figure 4), each row corresponds to a fixed shift *δ* between the distributions of the two populations, and *δ* gets larger from the top row to bottom row. In each row, the subfigure (a) shows the two population distributions, the subfigure (b) plots the powers for the balanced datasets with increasing *m* = *n*, and the subfigure (c) plots the powers for the unbalanced datasets with fixed *m* = 5 and increasing *n*. Our WMW-A test performs the best or similar with other test methods for almost all the datasets with either Normal distributions or Gamma distributions, either small shift *δ* or relatively big shift *δ*, either balanced case with the same *m* and *n* or unbalanced case with fixed *m* = 5 and increasing *n*. Especially, we have several observations from the two figures.

The WMW-A test performs the best when sample sizes are small, for both Normal and Gamma population distributions. This also reflects the robustness of the nonparametric test to the classes of population distributions. The t-test tends to have some advantages when sample sizes get relatively large for normal distributions, which is reasonable since t-test is based on the assumption of normal distributions.

The WMW-A test performs much better than the four comparison tests for small sample sizes for different *δ*s, either big or small. Large *δ* tends to generate higher powers, for all the five test methods, and the WMW-A test tends to perform similarly with other methods for the unbalanced case when sample sizes increase. Note that when *δ* = 0.5, the Welch test obtains higher power for Gamma distributions for with extremely unbalanced sample sizes (*m* = 5 and *n* is large), due to its advantages for inconsistent variances of the two samples. Overall, the WMW-A test could obtain higher power functions than other methods for almost all the cases.

The WMW-A test performs the best for most cases for either balanced or unbalanced datasets, and balanced datasets tend to generate more stable powers than the unbalanced datasets by all methods. When *m* or *n* increases in the balanced datasets, the powers increase stably. However, when *n* increases in the unbalanced datasets with fixed *m* = 5, the powers may not increase. It seems that the degree of unbalanceness influences the test powers. Interestingly, when the sample sizes are extremely small, for example, *m* = *n* = 2 for small delta (for example, *δ* = 0.5), the WMW-A test could obtain even higher power than *m* = *n* = 5. This implies the potential for the WMW-A test for extremely small sample sizes.

We also experimentally validated the theoretical asymptotic power in Theorem 3 when the sample sizes are large enough for asymptotic normal distributions. Figure 5 compares the theoretical powers approximated by Theorem 3 and the empirical powers calculated by permutation procedure, for data generated by Gamma distributions with different sample sizes. The results show that when *m* and *n* are both large, for either balanced case or unbalanced case, theoretical powers are similar to the empirical powers, while for small sample sizes, permutation powers tend to be higher.

**Figure 5.**
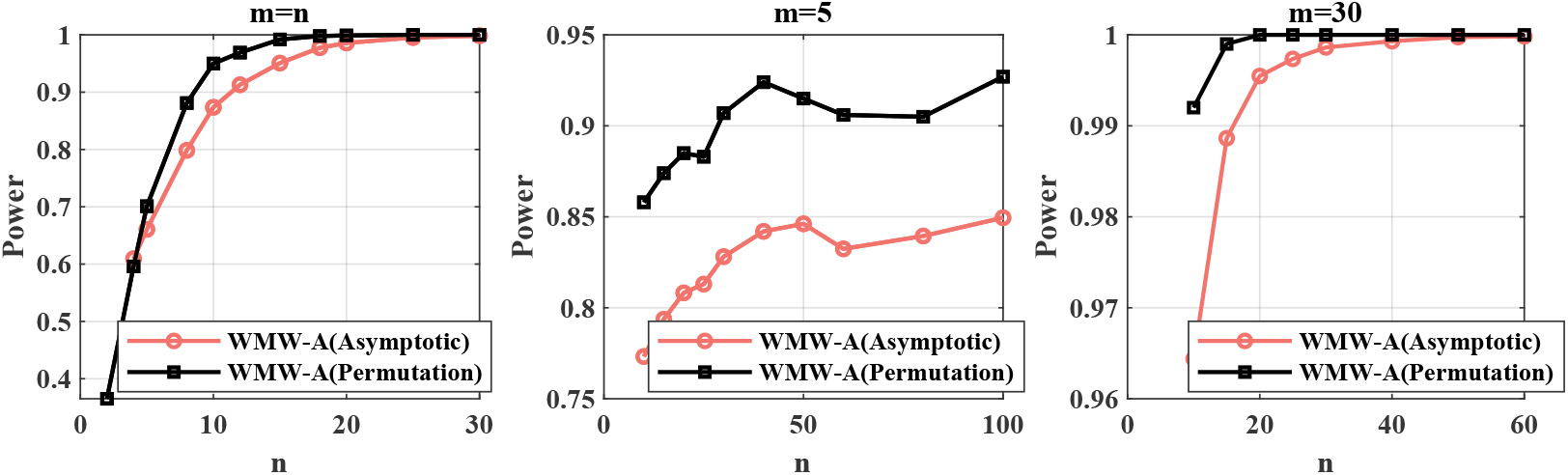
The theoretical asymptotic powers of WMW-A test approximated by 3, and the empirical powers calculated by permutation test, for both balanced and unbalanced cases. Datasets are generated from Gamma population distribution, where *α_x_* = 0.5, *α_y_* = 2, and *k* is chosen as 2.

We further checked the type I error rates of the different methods by simulating datasets with the same population distributions for *X* and *Y*. Normal distributions with *α_x_* = *α_y_* = 0, and Gamma distributions with *α_x_* = *α_y_* = 0.5 for both balanced case and unbalanced case(*m* = 5) were used to calculate type I error rates for different methods. Figure 6 reports the results. The two rows are corresponding to Normal and Gamma distributions, respectively, and the two columns are corresponding to the balanced case (*m* = *n*) and unbalanced case (*m* = 5), respectively. In each subfigure, the type I error rates with changed sample sizes were plotted. The WMW-A test tends to generate a little bit higher or similar type I error rates compared to the four compared test methods, as its acceptable price for the higher test powers. We also noticed that for the unbalanced case, the Welch test has much higher Type I error rates on the Gamma distribution, although it sometimes obtains higher power in the above experiments.

**Figure 6.**
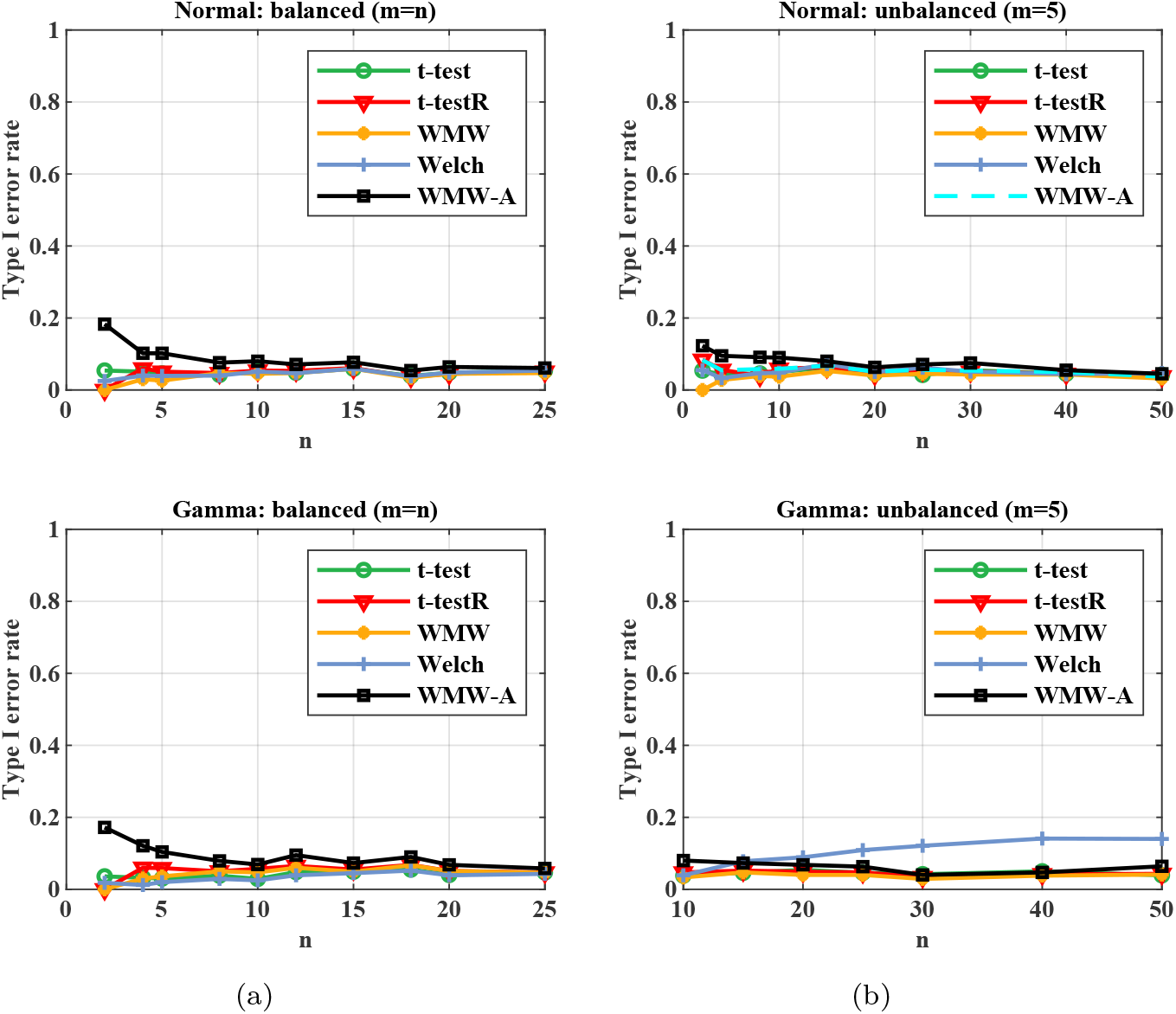
Type I error rates of t-test, t-testR, WMW test, Welch test and WMW-A test (*k* = 5) for datasets generated from Normal population distributions (the first row) and Gamma population distributions (the second row): (a).Type I error rates for the balanced case with increasing *m* = *n*; (b).Type I error rates for unbalanced case with fixed *m* = 5 and increasing *n*.

## Discussion about the auxiliary sample

In this section, we further discuss how the auxiliary sample influences the performance of WMW-A test, by changing the population distribution and the sample size of the auxiliary sample. We have three observations based on extensive experiments:

a. The larger sample size *k*(*m* + *n*) of auxiliary sample often implies higher power of the WMW-A test, but with a power limit when the auxiliary sample size increases.
b. When the unlabelled auxiliary data are available and *Z* follows a mixed distribution *F_Z_*(*x*) = *π_z_F_X_* (*x*) + (1 − *π_z_*)*F_Y_* (*x*), the WMW-A test is robust with regard to the parameter *π_z_*.
c. When the unlabelled auxiliary data is unavailable, they can be generated by an approximated Gaussian distribution (WMW-A-1G) or an approximated bimodal mixed Gaussian distribution (WMW-A-2G).

Firstly, large sample size of auxiliary *Z* tend to improve the power of WMW-A test, especially when the sample sizes of *X* and *Y* are small. We set the sample size of auxiliary sample as *l* = *k*(*m* + *n*), and checked the performance of the WMW-A test with fixed *π_z_* = *π* and different *k* on simulation datasets. Figure 7 shows that the WMW-A test with larger *k* tends to obtain higher powers for two-small-sample test. An obvious disadvantage of large sample size of *Z* is the high computation complexity. One may need to balance the performance and the computation.

**Figure 7.**
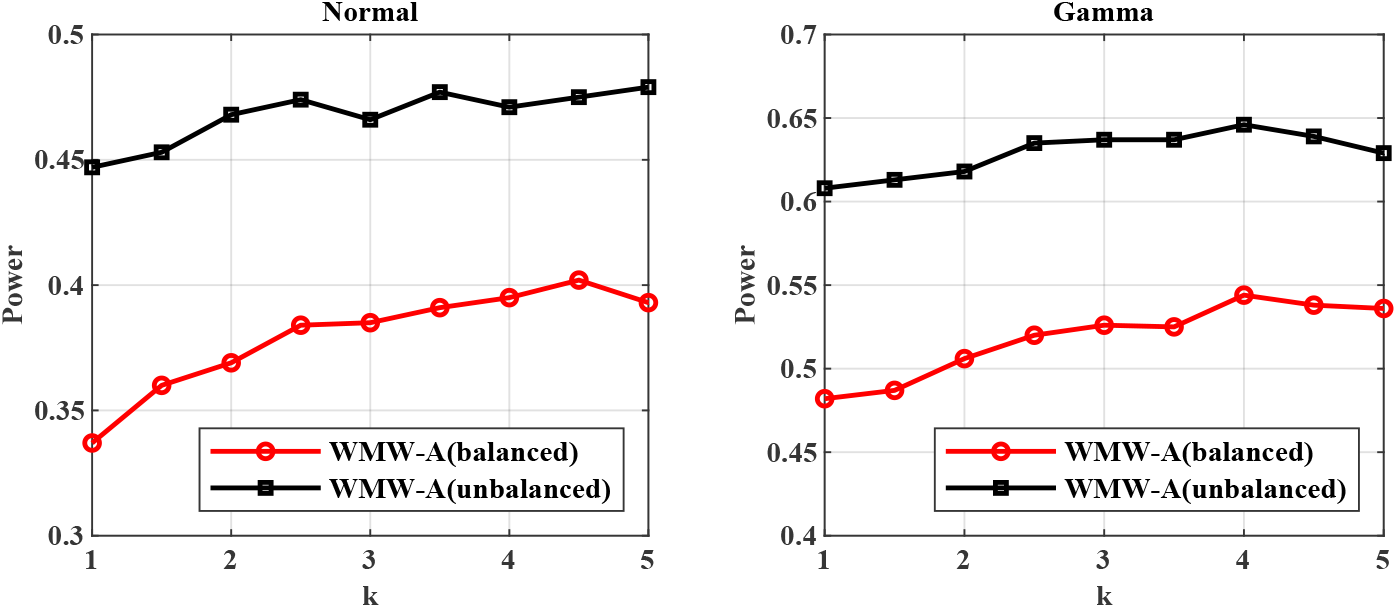
Powers of WMW-A test for datasets generated from Normal and Gamma population distributions with increasing *k* under balanced case (*m* = *n* = 5) and unbalanced case (*m* = 5, *n* = 10), where *α_x_* = 0.5, *α_y_* = 1.5, *π_z_* = *π*.

Secondly, when there are large amount of available unlabelled auxiliary data, which follows a mixed distribution of *X* and *Y*, the WMW-A test is robust to the parameter *π_z_* of the mixed distribution. It implies that the proportions of case and control in the unlabelled data do not have much influence on the performance of the WMW-A test. Note that *π* is a parameter for simulating the case and control observations, and it represents the proportion of the pooled *X* − *Y* sample which are from *X*. *π_z_* is a parameter for selecting *Z*-sample in the WMW-A test, and it represents the proportion of *Z* which are sampled from the population *X*. We checked the performance of the WMW-A test with different *π_z_* on datasets generated by different *π*. We fixed the total sample size *m* + *n* = 30, and generated case and control data by Gamma distributions with *δ* = 0.5 and *δ* = 1, respectively, with different proportion *π* from 0.1 to 0.9. To perform the WMW-A test, we fixed *k* = 1 and varied the parameter *π_z_* from 0 to 1. Powers were computed and reported in Figure 8. The results showed that the WMW-A test is robust with regard to the parameter *π_z_*, since the results did not change much when *π_z_* changed.

**Figure 8.**
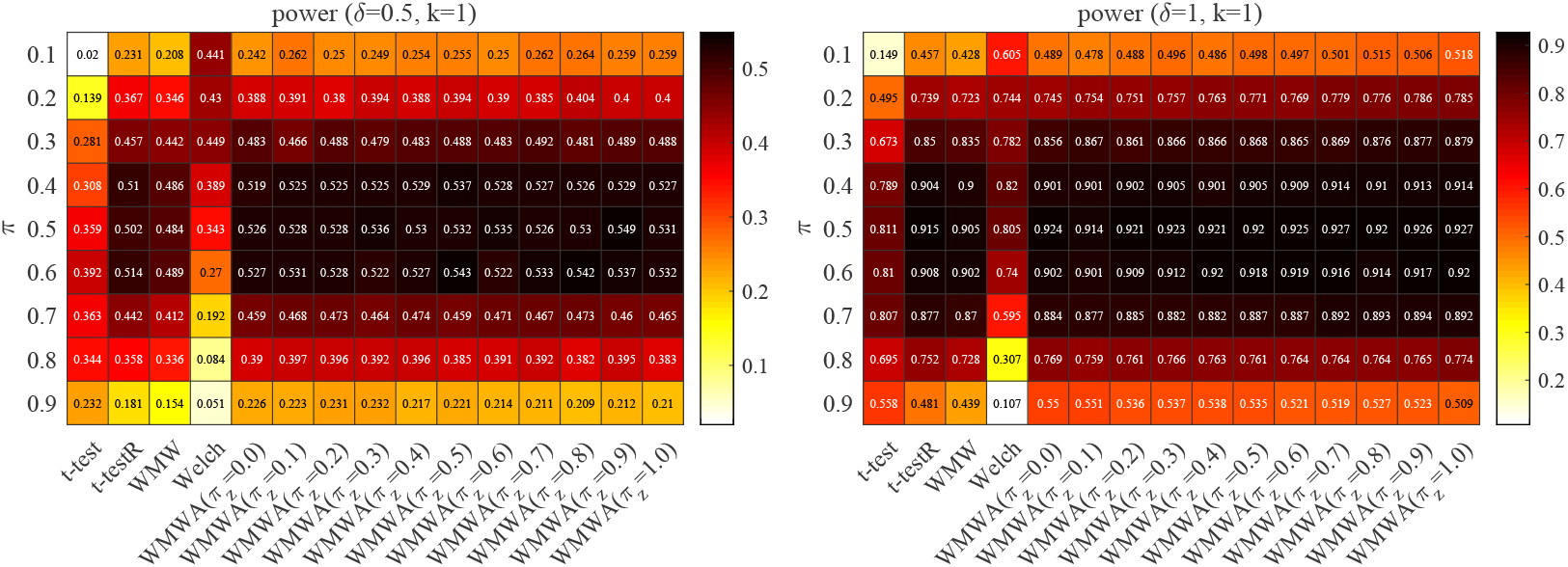
Powers of t-test, t-testR, WMW test, Welch test and WMW-A test for datasets generated from Gamma distributions. WMW-A test with different *π_z_* on datasets generated by different *π*, for *δ* = 0.5 and *δ* = 1, respectively, where *π* = {0.1, · · ·, 0.9}, *π_z_* = {0, · · ·, 1} and *m* + *n* = 30, *k* = 1.

Finally, when the unlabelled auxiliary data is unavailable in applications, the generated auxiliary data could also be used in WMW-A test. We proposed two ways to generate auxiliary samples and perform WMW-A test, the procedures are called as WMW-A-1G and WMW-A-2G, respectively. The WMW-A-1G procedure samples auxiliary data from a single normal distribution whose mean and variance are approximated by the case and control observations together. The WMW-A-2G procedure samples auxiliary data from a mixed Gaussian distribution by the MH step [13], whose distribution parameters are approximated by the case and control observations. To aviod confusion, WMW-A test represents the test with available unlabelled auxiliary data. The power and type I error rates of the WMW-A-1G and WMW-A-2G are both reported in Table 3. We can see that the powers of our WMW-A, WMW-A-1G and WMW-A-2G test are higher than the compared methods in all of the cases, and the WMW-A-1G procedure and WMW-A-2G procedure with generated auxiliary data have comparative performance with the WMW-A test with available auxiliary data. Furthermore, the WMW-A-2G procedure tends to outperform the WMW-A-1G procedure.

**Table 3.** Powers and Type I error rates of t-test, t-testR, WMW test, Welch test and WMW-A test, WMW-A-1G test, WMW-A-2G test with *k* = 5 for datasets generated from Normal and Gamma population distributions under the balanced case (*m* = *n* = 5).

|  | distribution | t-test | t-testR | WMW | Welch | WMW-A | WMW-A-1G | WMW-A-2G |
| --- | --- | --- | --- | --- | --- | --- | --- | --- |
| Power | Normal | 0.285 | 0.290 | 0.220 | 0.267 | 0.395 | 0.390 | 0.412 |
|  | Gamma | 0.311 | 0.413 | 0.313 | 0.24 | 0.538 | 0.517 | 0.525 |
| Type I error rate | Normal | 0.056 | 0.055 | 0.036 | 0.052 | 0.098 | 0.100 | 0.137 |
|  | Gamma | 0.028 | 0.056 | 0.032 | 0.017 | 0.104 | 0.138 | 0.151 |

Overall, the WMW-A test is robust to the selection of *Z*, specifically including the sample size, the proportion *π_z_* and the population distributions, and thus it could be widely used in real applications.

## 1 Real applications on gene expression data

In this section, we evaluate the WMW-A method for two-small-sample test by real biological gene expression datasets. We collect datasets from the GEO database, which is a gene expression database created and maintained by the National Biotechnology Information Center NCBI, and contains a large number of high-throughput gene expression datasets submitted by research institutions around the world. We evaluate the WMW-A test on the following three gene expression datasets.

a. The breast cancer (BC) dataset is accessible through GEO Series accession number GSE86374. The experiment used Affymetrix microarray analysis to study potential breast cancer diagnostic markers and further analyzed the differential genes between subtypes of breast cancer. The dataset includes 124 breast cancer samples and 35 healthy samples with 33297 genes.
b. The colorectal cancer (CRC) dataset is accessible through GEO Series accession number GSE83889. This experiment studied the gene expression data for 47323 genes from 101 colorectal cancer tissues and 35 matched non-tumor colon mucosal tissues, and the marker genes are identified after subtype identification on the colorectal cancer samples.
c. The Peripheral T-cell Lymphoma (PTCL) dataset is accessible through GEO Series accession number GSE132550. This experiment studied the molecular events of peripheral T-cell lymphomas, not otherwise specified (PTCL/NOS), and analyzed gene expression of PTCL/NOS cases and normal T cells by DASL array. The PTCL dataset includes 18023 genes for 80 patients and 10 healthy normal samples.

The summary of the three datasets is shown in Table 4.

**Table 4.**
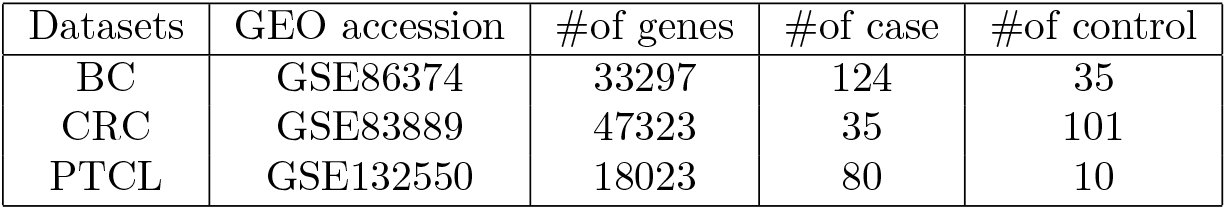
Summary of the datasets including GEO accession, the number of genes, and the sample size of case and control.

To evaluate the performance of WMW-A test, We perform WMW-A test and four comparison test (t-test, t-testR,WMW test and Welch test) on the three datasets by selecting *m* cases and *n* controls, where (*m, n*) are chosen as (5, 5), (5, 10), (5, 15), (5, 20), (10, 10), (15, 15), respectively. For WMW-A test, we take both available unlabelled auxiliary data (WMW-A), and generated auxiliary data (WMW-A-1G and WMW-A-2G). The parameter setting of auxiliary is set as *k* = 2.

To approximate the test powers and type I error rates for each method, we use the ground truth differential genes obtained by the analysis tool of GEO2R [9], [20], which is included in the GEO database and used to analyze the statistical information of each gene, such as *p* value, etc. Then the differential genes are filtered according to the commonly used filtering thresholds: *p* < 0.01 and |log_2_ Fold Change| ≥ 1. We report the average power and type I error rate of the WMW-A test, where the power is the proportion of the correctly found differential genes. We performed the FDR correction by the BH method [2] for the multiple test results. The results of the three datasets are shown in Table 5 and 6. As shown in Table 5, for all the three datasets, the WMW-A test could obtain higher powers than the compared methods. The WMW-A test with either the available auxiliary data (WMW-A) or generated auxiliary data (WMW-A-1G or WMW-A-2G) could siginificantly improve the test power. The WMW-A-1G and WMW-A-2G test with generated auxiliary data sometimes achieve even higher power than the WMW-A test, which verify the effectiveness of the generated auxiliary data. Although the type I error rates of our method in Table 6 are higher, they are mostly controlled in an acceptable region in real applications. Overall, these results verify the effectiveness of the WMW-A test for small sample sizes.

**Table 5.** Powers of t-test, t-testR, WMW test, Welch test and WMW-A test, WMW-A-1G test, WMW-A-2G test for the three gene expression datasets under significance level *α* = 0.05 based on the ground truth from GEO2R.

|  | Methods | (5,5) | (5,10) | (5,15) | (5,20) | (10,10) | (15,15) |
| --- | --- | --- | --- | --- | --- | --- | --- |
| BC | t-test | 0.059 | 0.267 | 0.297 | 0.459 | 0.462 | 0.806 |
|  | t-testR | 0.199 | 0.223 | 0.297 | 0.368 | 0.504 | 0.816 |
|  | WMW | 0.000 | 0.156 | 0.033 | 0.081 | 0.250 | 0.772 |
|  | Welch | 0.023 | 0.116 | 0.210 | 0.294 | 0.381 | 0.792 |
|  | WMW-A | 0.428 | 0.563 | 0.549 | 0.630 | 0.703 | 0.873 |
|  | WMW-A-1G | 0.339 | 0.485 | 0.578 | 0.566 | 0.707 | 0.867 |
|  | WMW-A-2G | 0.243 | 0.400 | 0.447 | 0.546 | 0.647 | 0.813 |
| CRC | t-test | 0.501 | 0.719 | 0.782 | 0.817 | 0.931 | 0.981 |
|  | t-testR | 0.698 | 0.790 | 0.840 | 0.876 | 0.928 | 0.977 |
|  | WMW | 0.000 | 0.700 | 0.598 | 0.775 | 0.899 | 0.974 |
|  | Welch | 0.294 | 0.810 | 0.876 | 0.908 | 0.908 | 0.978 |
|  | WMW-A | 0.775 | 0.875 | 0.903 | 0.923 | 0.951 | 0.983 |
|  | WMW-A-1G | 0.831 | 0.910 | 0.933 | 0.945 | 0.962 | 0.988 |
|  | WMW-A-2G | 0.703 | 0.841 | 0.879 | 0.895 | 0.948 | 0.981 |
| PTCL | t-test | 0.440 | 0.588 | 0.690 | 0.725 | 0.917 | — |
|  | t-testR | 0.668 | 0.779 | 0.841 | 0.864 | 0.927 | — |
|  | WMW | 0.486 | 0.736 | 0.789 | 0.833 | 0.911 | — |
|  | Welch | 0.232 | 0.784 | 0.869 | 0.886 | 0.898 | — |
|  | WMW-A | 0.671 | 0.790 | 0.836 | 0.855 | 0.930 | — |
|  | WMW-A-1G | 0.563 | 0.714 | 0.759 | 0.784 | 0.905 | — |
|  | WMW-A-2G | 0.715 | 0.829 | 0.871 | 0.887 | 0.946 | — |

**Table 6.**
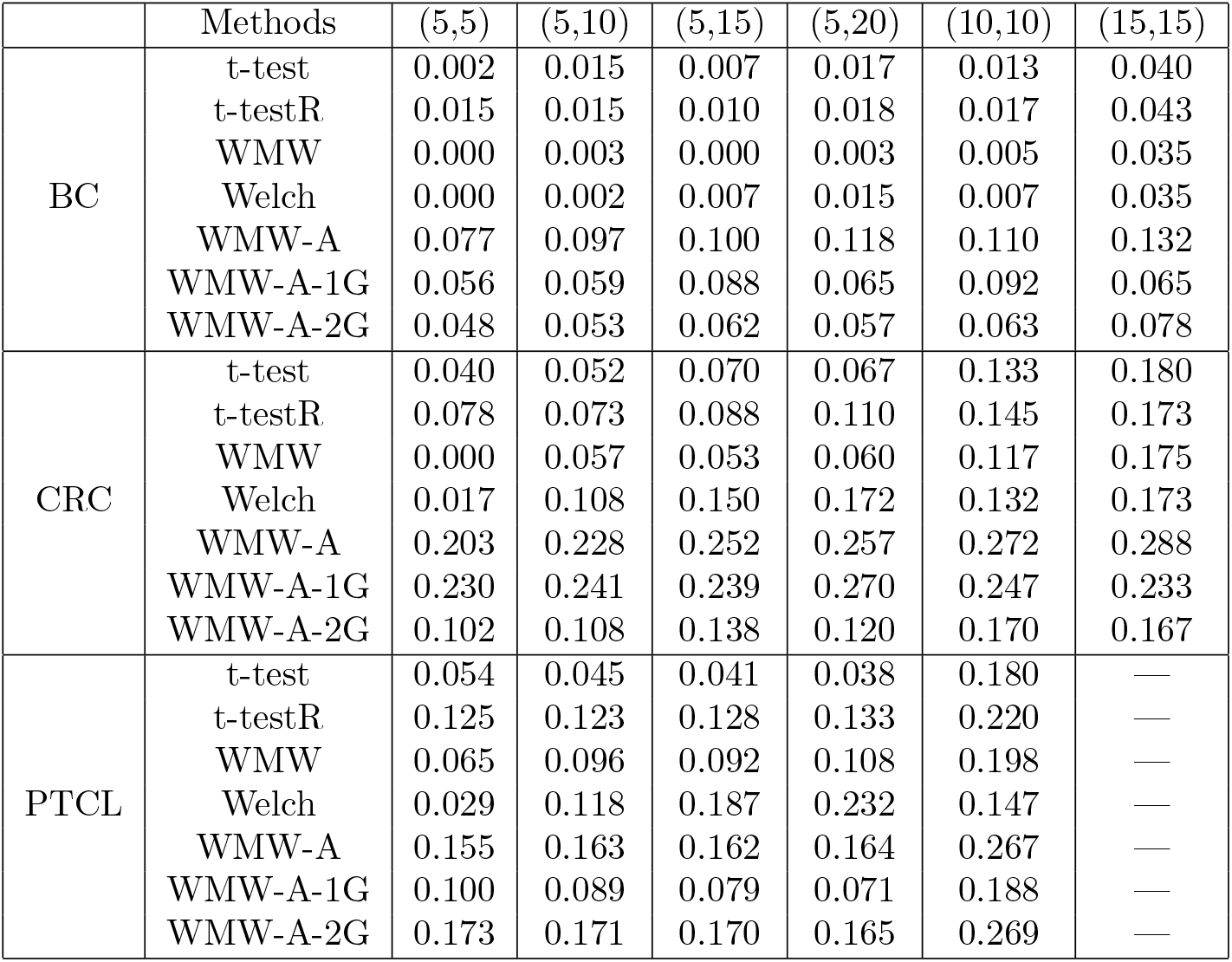
Type I error rates of t-test, t-testR, WMW test, Welch test and WMW-A test, WMW-A-1G test, WMW-A-2G test for the three gene expression datasets under significance level *α* = 0.05 based on the ground truth from GEO2R.

## Conclusion

Small sample size is an interesting and challenging problem in the big data era. Much effort has been done to deal with the small sample problem. However, few work considers two-sample test in the case of small sample size, which exists in many biological scenarios. Here, by small sample size, we mean less than 20, or even 10. In this work, we proposed a novel nonparametric two-sample test, the WMW-A test, for small sample sizes by introducing an auxiliary sample. The main idea is to first pool the control, case and auxiliary samples together, and then to rank the *X*-sample and *Y* -sample. The WMW-A statistic is constructed by the gap between their average ranks. Large WMW-A statistic implies far distance between the *X* and *Y* samples among the pooled samples. We theoretically analysed the properties of the WMW-A statistics, such as the symmetry, its means and variances under null hypothesis and alternative hypothesis. We further theoretically analysed the test power of the WMW-A test. The permutation procedure was then given in the WMW-A test algorithm.

We evaluated the WMW-A test by both simulation and real biological datasets, and the experimental results show better performance of the WMW-A test than traditional tests for small sample sizes. We further checked the robustness of the selection of *Z*-sample. The results show that the WMW-A test is robust to the selection of auxiliary distributions, and it could perform well by either available unlabelled auxiliary data or generated auxiliary data. This guarantees that the WMW-A test can be easily implied to real datasets. Our experimental results on the biological gene expression datasets further validated the effectiveness of the WMW-A test.

Small sample problem is a very difficult problem in statistics, since most statistical methods are only applicable for large sample sizes. It is a brand new idea to introduce an auxiliary sample to improve the test power with an acceptable type I error rate. Future work might consider the small sample unbalanced problem and how to further control type I error rates to improve the test performance.

## Acknowledgments

We thank just about everybody.

## Footnotes

1 The source code of onelearn is available at https://github.com/LiminLi-xjtu/WMW-A

## References

1. Z. Bai and H. Saranadasa. Effect of high dimension: by an example of a two sample problem. Statistica Sinica, pages 311–329, 1996.

2. Y. Benjamini and Y. Hochberg. Controlling the false discovery rate: a practical and powerful approach to multiple testing. Journal of the Royal statistical society: series B (Methodological), 57(1):289–300, 1995.

3. Y. Cao, W. Lin, and H. Li. Two-sample tests of high-dimensional means for compositional data. Biometrika, 105(1):115–132, 2017.

4. H. Chen, X. Chen, and Y. Su. A weighted edge-count two-sample test for multivariate and object data. Journal of the American Statistical Association, 113(523):1146–1155, 2018.

5. H. Chen and J. H. Friedman. A new graph-based two-sample test for multivariate and object data. Journal of the American statistical association, 112(517):397–409, 2017.

6. S. X. Chen, Y.-L. Qin, et al. A two-sample test for high-dimensional data with applications to gene-set testing. The Annals of Statistics, 38(2):808–835, 2010.

7. W. J. Conover and R. L. Iman. Rank transformations as a bridge between parametric and nonparametric statistics. The American Statistician, 1981.

8. D. A. Darling. The kolmogorov-smirnov, cramer-von mises tests. The Annals of Mathematical Statistics, 28(4):823–838, 1957.

9. S. Davis and P. Meltzer. Geoquery: a bridge between the gene expression omnibus (geo) and bioconductor. Bioinformatics, 14:1846–1847, 2007.

10. Y. Fong, Y. Huang, M. P. Lemos, and M. J. Mcelrath. Rank-based two-sample tests for paired data with missing values. Biostatistics, 19(3):281–294, 2018.

11. J. H. Friedman and L. C. Rafsky. Multivariate generalizations of the wald-wolfowitz and smirnov two-sample tests. The Annals of Statistics, pages 697–717, 1979.

12. K. B. Gregory, R. J. Carroll, V. Baladandayuthapani, and S. N. Lahiri. A two-sample test for equality of means in high dimension. Journal of the American Statistical Association, 110(510):837–849, 2015.

13. W. K. Hastings. Monte carlo sampling methods using markov chains and their application. Biometrika, 57(1), 1970.

14. S. Hediger, L. Michel, and J. Näf. On the use of random forest for two-sample testing. arXiv preprint arXiv:1903.06287, 2019.

15. I. Kim, A. B. Lee, J. Lei, et al. Global and local two-sample tests via regression. Electronic Journal of Statistics, 13(2):5253–5305, 2019.

16. H. B. Mann and D. R. Whitney. On a test of whether one of two random variables is stochastically larger than the other. The annals of mathematical statistics, pages 50–60, 1947.

17. E. Olivetti, S. Greiner, and P. Avesani. Statistical independence for the evaluation of classifier-based diagnosis. Brain informatics, 2(1):13–19, 2015.

18. R. Peto and J. Peto. Asymptotically efficient rank invariant test procedures. Journal of the Royal Statistical Society: Series A (General), 135(2):185–198, 1972.

19. P. Philonenko and S. Postovalov. The new robust two-sample test for randomly right-censored data. Journal of Statistical Computation and Simulation, 89(8):1357–1375, 2019.

20. M. E. Ritchie, B. Phipson, D. Wu, Y. Hu, C. W. Law, W. Shi, and G. K. Smyth. limma powers differential expression analyses for RNA-sequencing and microarray studies. Nucleic Acids Research, 43(7):e47, 2015.

21. P. R. Rosenbaum. An exact distribution-free test comparing two multivariate distributions based on adjacency. Journal of the Royal Statistical Society: Series B (Statistical Methodology), 67(4):515–530, 2005.

22. J. D. Rosenblatt, Y. Benjamini, R. Gilron, R. Mukamel, and J. J. Goeman. Better-than-chance classification for signal detection. arXiv preprint arXiv:1608.08873, 2016.

23. S.-i. Tsukada. High dimensional two-sample test based on the inter-point distance. Computational Statistics, 34(2):599–615, 2019.

24. A. W. Van der Vaart. Asymptotic statistics, volume 3. Cambridge university press, 2000.

25. A. Wald and J. Wolfowitz. On a test wether two samples are from the same distribution. Ann. Math. Stat, 11:147–162, 1940.

26. B. L. Welch. The significance of the difference between two means when the population variances are unequal. Biometrika, 29, 1938.

27. F. Wilcoxon. Individual comparisons by ranking methods. Biometrics, 1(6):80–83, 1945.

28. Y. Wu, M. G. Genton, and L. A. Stefanski. A multivariate two-sample mean test for small sample size and missing data. Biometrics, 62(3):877–885, 2006.

29. K. Xue and F. Yao. Distribution and correlation free two-sample test of high-dimensional means. arXiv preprint arXiv:1904.07416, 2019.

30. J.-T. Zhang, J. Guo, B. Zhou, and M.-Y. Cheng. A simple two-sample test in high dimensions based on l 2-norm. Journal of the American Statistical Association, pages 1–42, 2019.

